# SPARTA: Simple Program for Automated reference-based bacterial RNA-seq Transcriptome Analysis

**DOI:** 10.1101/021915

**Authors:** Benjamin K. Johnson, Matthew B. Scholz, Tracy K. Teal, Robert B. Abramovitch

**Affiliations:** Department of Microbiology and Molecular Genetics, Michigan State University, East Lansing, Michigan, 48824, USA; Institute for Cyber-Enabled Research, Michigan State University, East Lansing, Michigan, 48824, USA; Data Carpentry

## Abstract

**Summary:** SPARTA is a reference-based bacterial RNA-seq analysis workflow application for single-end Illumina reads. SPARTA is turnkey software that simplifies the process of analyzing RNA-seq data sets, making bacterial RNA-seq analysis a routine process that can be undertaken on a personal computer or in the classroom. The easy-to-install, complete workflow processes whole transcriptome shotgun sequencing data files by trimming reads and removing adapters, mapping reads to a reference, counting gene features, calculating differential gene expression, and, importantly, checking for potential batch effects within the data set. SPARTA outputs quality analysis reports, gene feature counts and differential gene expression tables and scatterplots. The workflow is implemented in Python for file management and sequential execution of each analysis step and is available for Mac OS X, Microsoft Windows, and Linux. To promote the use of SPARTA as a teaching platform, a web-based tutorial is available explaining how RNA-seq data are processed and analyzed by the software.

**Availability and Implementation:** Tutorial and workflow can be found at sparta.readthedocs.org. Teaching materials are located at sparta-teaching.readthedocs.org. Source code can be downloaded at www.github.com/abramovitchMSU/, implemented in Python and supported on Mac OS X, Linux, and MS Windows.

**Contact:** Robert B. Abramovitch

**Supplemental Information:** Supplementary data are available at *Bioinformatics* online.

## 1 INTRODUCTION

One of the most common applications of RNA sequencing (RNA-seq) is to identify differentially expressed genes under differing experimental conditions. Before biological insights can be gained, one must process and analyze the large datasets generated from each sequencing experiment. Each sample contains millions of reads that must be trimmed and assessed for read quality, mapped back to a reference genome (or assembled *de novo* in the absence of a reference), counted for transcript abundance, and tested for differential gene expression. Many computational analysis tools have been developed specifically to work with RNA-seq data; however, a single tool is often not suitable and requires several different applications assembled into a workflow. This task can be complicated as both the tool choice and input and output file formats for a given tool need to be considered and potentially modified to meet the requirements for the subsequent analysis step. Several RNA-seq analysis workflows exist, however, most are designed for eukaryotic organisms (D’Antonio, et al., 2015; Golosova, et al., 2014; Goncalves, et al., 2011; Habegger, et al., 2011; Kalari, et al., 2014; McClure, et al., 2013; Michalovova, et al., 2015; Qi, et al., 2010; Tjaden, 2015; Wang, et al., 2011). The goal of this work is to assemble several open-source computational tools to deliver a complete, accessible, and easy-to-use reference-based bacterial RNA-seq analysis workflow that is amenable to both the research laboratory and undergraduate classroom.

## 2 FEATURES AND FUNCTIONS

The SPARTA workflow (Figure 1) is implemented utilizing Python for file input/output management and tool execution, combining several open-source computational tools. The SPARTA workflow analyzes data by: conducting read trimming and adapter removal with Trimmomatic (Bolger, et al., 2014); performing quality analysis of the data sets with FastQC (Anders, 2010); mapping the reads to the reference with Bowtie (Langmead, et al., 2009); counting transcript or gene feature abundance with HTSeq (Anders, et al., 2015); and, analyzing differential gene expression with edgeR (McCarthy, et al., 2012; Robinson, et al., 2010; Robinson and Oshlack, 2010). Within the differential gene expression analysis step, batch effects can be detected and the user is warned that potentially unintended variables need to be considered. If left unaccounted for, batch effects can significantly skew the results of the data analysis, leading to inappropriate experimental conclusions (Leek, et al., 2010). Following analysis, SPARTA outputs quality analysis reports, gene feature counts and differential gene expression tables and scatterplots.

**Figure 1.**
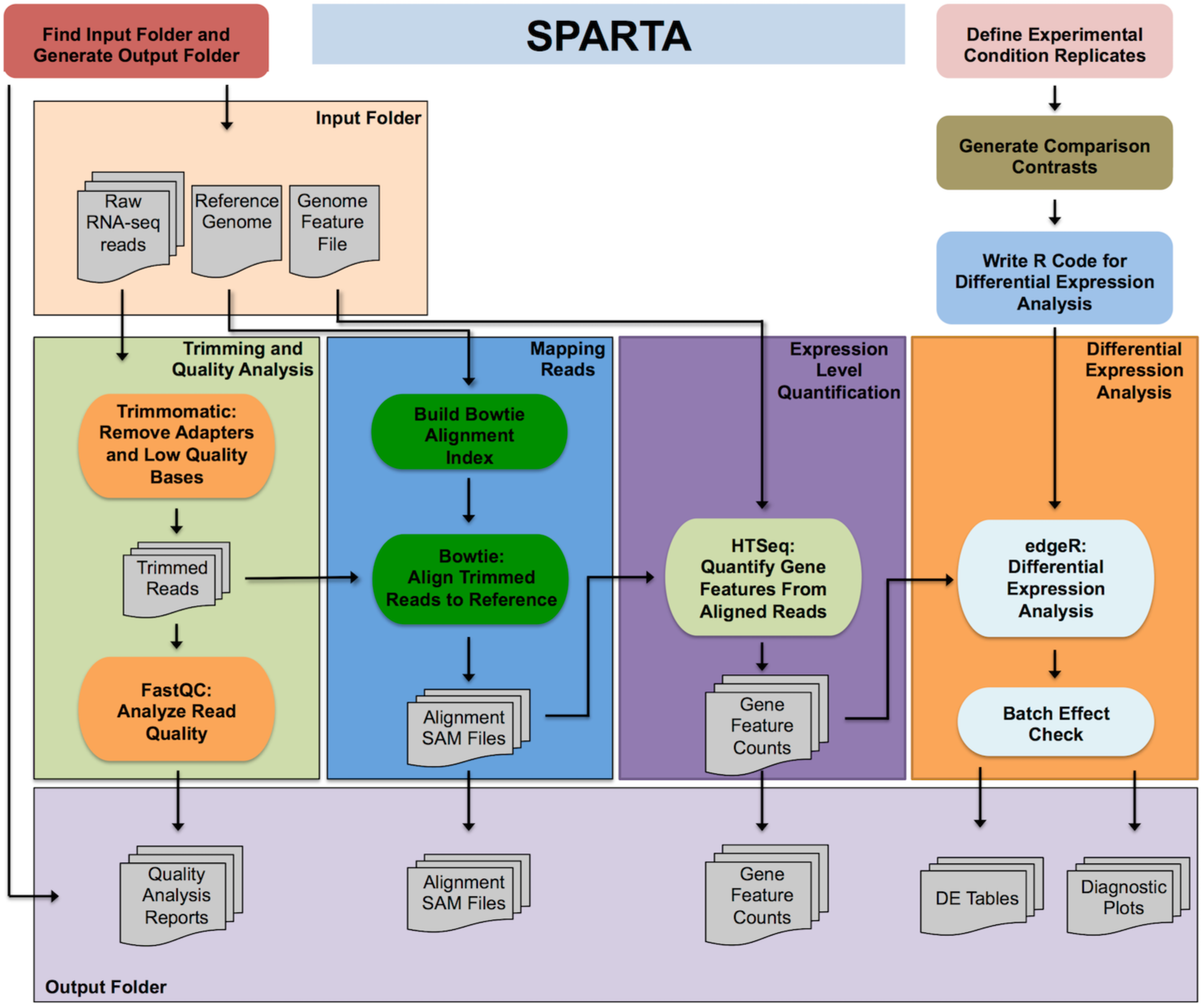
SPARTA workflow diagram

SPARTA requires Python 2, NumPy (a Python library for numerical analyses), Java and R. Once Python is installed, the user initializes SPARTA, which then checks for the necessary dependencies at runtime. If any of these dependencies are not met, SPARTA informs the user of the missing components. To reduce complex software installation, SPARTA is distributed with the required software and an online tutorial (sparta.readthedocs.org) guides the user through installation and data analysis procedures for each operating system platform. The workflow maintains analytic flexibility for specific use cases by allowing the user to tailor the options utilized for each analysis step, but can proceed without requiring option specification. Further, SPARTA will write the necessary R commands at runtime and will generate the appropriate contrasts to test all possible comparisons between user defined experimental conditions. Using a previously published data set (Baker, et al., 2014), SPARTA was capable of analyzing 4 experimental conditions containing 8 samples with approximately 30 million reads per sample in 4 hours on an off-the-shelf iMac computer (8 GB RAM, Intel i5 2.7GHz quad-core processor). SPARTA can also be implemented in high performance computing environments utilizing the non-interactive mode functionality (Supplemental material).

As NGS technologies and applications continue to permeate life science research, undergraduate education must include the use of contemporary sequencing techniques to address biological questions. However, despite the rapid increase in data intensive experimental biology, undergraduates receiving a life sciences degree are often not exposed to the tools and basic computational skills required to study NGS data sets. To address this shortcoming, we have developed an online tutorial to guide students through the RNA-seq analysis process (sparta-teaching.readthedocs.org). The SPARTA teaching tool was integrated into a senior level genomics course and successfully engaged students in the theory and application of RNA-seq data analysis.

In summary, RNA-seq transcriptional profiling is becoming increasingly routine, and there is a demand for applications such as SPARTA that enable stand-alone workflows. SPARTA represents an easy-to-use, platform independent analysis workflow for reference-based bacterial RNA-seq amenable to the research laboratory and classroom.

## ACKNOWLEDGEMENTS

This project was supported by grants to RBA from the NIH (R21AI105867) and the Bill & Melinda Gates Foundation (OPP1119065).

